# BioWorldModel: a single architecture predicts phenotype from genotype across four kingdoms of life

**DOI:** 10.64898/2026.03.27.714912

**Authors:** Khasim Hussain Baji Shaik, Ankur Sahu

## Abstract

The same genome produces different phenotypes in different conditions—yet predictive models encode genotype once and treat each trait independently. Here we show that representing phenotype generation as a dynamic biological process transforms predictive accuracy across bacteria, fungi, animals and plants.

BioWorldModel learns how organisms interpret their genome: frozen gene embeddings (species context) modulated by individual variation pass through four biological process layers (regulation → expression → pathway → cellular) that respond to environment and time. A state-conditioned attention mechanism rereads this dynamic representation, predicting full multivariate trait distributions.

Without modification, the architecture achieves mean correlation *r* = 0.678 on 214 bacterial growth traits (207% better than ridge regression), *r* = 0.915 on 35 yeast fitness traits (167% better), *r* = 0.499 on 199 fly phenotypes in small-sample regime (760% better), and *r* = 0.995 on 36 rice traits (49% better). Ablations confirm that modeling biological process—not model size—drives performance. When neural architectures represent how biology generates phenotype rather than merely associating genotype with outcome, they capture what static methods miss.

## 1 Introduction

A bacterial strain resistant to one antibiotic may be sensitive to another. A rice variety high-yielding in wet conditions may fail under drought. The same genome produces different phenotypes depending on cellular state, environment and developmental time ^1;2^. This is not measurement noise—it is biology. Genes are not decoded; they are *interpreted*. Interpretation depends on context. Phenotype emerges from this dynamic reading, not from static genomic features.

Standard genomic prediction ignores this. Ridge regression, random forests, and deep networks ^3–6^ encode genotype once, train one model per trait, and predict phenotype in a single forward pass. These methods have been useful—they power plant and animal breeding worldwide ^7^.

But their assumptions impose limits. If genotype is encoded once, it cannot express context-dependent function. If traits are predicted independently, the model cannot learn pleiotropy. If biological state is absent, prediction becomes pattern matching rather than process modeling.

We reasoned that prediction accuracy would improve if architectures represented *how* organisms generate phenotype. The question is not whether neural networks can fit genotype-phenotype data—they can. The question is whether an architecture designed around biological interpretation outperforms one designed around statistical association.

Here we introduce **BioWorldModel**: a system that separates species-level gene function from individual variation, transforms genomic representation through explicit biological process layers, conditionally rereads that representation as organismal state changes, and forecasts full multivariate phenotype distributions.

The critical test is generalization. If biological structure drives performance, the same architecture should work across vastly different organisms without retuning. We evaluate this claim on four datasets spanning bacteria (*Escherichia coli*), fungi (*Saccharomyces cerevisiae*), animals (*Drosophila melanogaster*), and plants (*Oryza sativa*)—205 to 971 individuals, 35 to 214 traits, 3,221 to 83,343 genetic markers. Hyperparameters are fixed. Architecture is identical. Only input data changes.

The result is unambiguous: biological structure wins. BioWorldModel outperforms per-trait baselines by 49–760% depending on organism and sample size. Ablations confirm that this advantage comes from modeling biological process—BioProcessStack layers, conditional genome reading, multivariate output—not from parameter count. When you model the generative process, you capture signal that association methods miss.

## 2 Results

### 2.1 BioWorldModel represents phenotype generation as biological interpretation

The architecture instantiates one biological principle: genomes are reread conditionally, not decoded once (Fig. 1). The same DNA sequence produces different molecular states depending on cellular context—environment, time, physiological condition. Four architectural innovations implement this.

**Figure 1:**
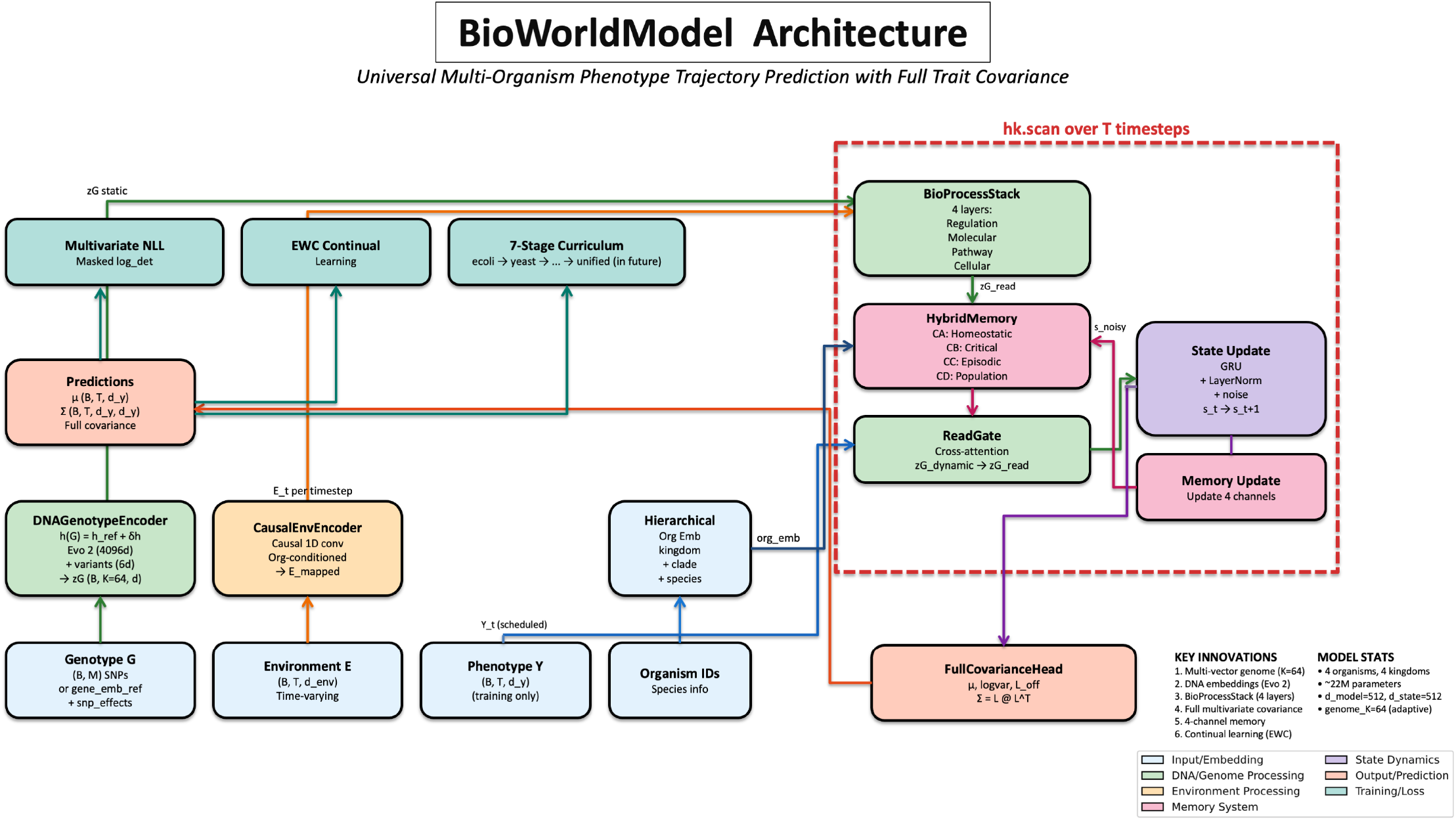
BioWorldModel architecture: same genome, different conditions, different biology. (**a**) Gene embeddings from frozen Evo 2 (species reference) are modulated by per-individual variant summaries. Attention pooling produces *K* = 64 genomic vectors: *h*(individual) = *h*_ref_(species) + *δh*(variants). (**b**) Four biological process layers (Regulation→ Expression→ Pathway→ Cellular) transform static **z**_*G*_ into dynamic 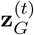, modulated by environment *e*_*t*_ and time *t*. Each layer: *z*_out_ = *z*_in_ + gate(*e*_*t*_, *t*) transform(*z*_in_). Same genome, different conditions → different representation. (**c**) ReadGate: a query from [environment, state, memory] attends over the 64 dynamic genomic vectors, retrieving condition-relevant signal. (**d**) Four biological memory channels (homeostatic, developmental, episodic, population) are gated and combined. GRU updates state *s*_*t*_. Multivariate head predicts *µ*_*t*_ and full covariance ∑_*t*_ = *LL*^*T*^ . **Core principle**: DNA is the book. Environment decides which chapter is read. Time decides the page. The model learns *how* biology interprets genotype.

#### Innovation 1: Frozen evolutionary context + learned individual variation

Standard genomic prediction encodes each individual’s genotype de novo. We factorize gene representation into two components:

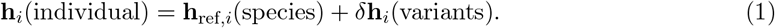

The reference **h**_ref,*i*_ is a 4,096-dimensional embedding from Evo 2 7B ^8^, frozen during training. This captures *what the gene does*: evolutionary constraints, functional neighborhoods, regulatory logic—knowledge amortized across millions of sequences. The modulation *δ***h**_*i*_ is learned from six per-gene features computed from this individual’s SNPs: heterozygous count, homozygous-alternate count, missing count, total SNPs, mean dosage, dosage variance. A gated projection determines how variants perturb the reference:

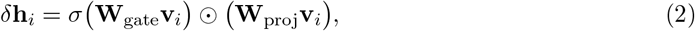

where **v**_*i*_ ∈ ℝ^6^ are the variant features. This separates *what a gene does* (universal, frozen) from *what this individual has* (population-specific, learned). Multi-vector attention pooling with 64 organism-conditioned queries ^9^ compresses thousands of genes into **z**_*G*_ ∈ ℝ^64×512^—a compact, interpretable genome representation.

#### Innovation 2: Environment-modulated biological process layers

Static genomes do not directly produce phenotype. Biology interprets DNA through hierarchical molecular processes that respond to conditions. We model this explicitly via BioProcessStack—four layers representing **Regulation** (which genes are accessible), **Expression** (what molecules exist), **Pathway** (what processes are active), and **Cellular** (what the organism is doing). Each layer transforms the genome representation via environment-gated residual connections:

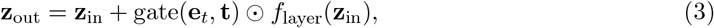

where gate(**e**_*t*_, **t**) = *σ*(**W**_gate_[**e**_*t*_; **t**_emb_]) is a sigmoid modulation signal from environment and time. The same genotype **z**_*G*_ yields different molecular state 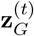 under drought vs. flood, starvation vs. feast—*exactly as biology works*. This is not a generic residual network; it is a mechanistic model of conditional gene expression.

#### Innovation 3: State-conditioned genome reading

Which genes matter depends on where the organism is in biological state space. High glucose activates glycolysis genes in yeast; ethanol activates respiration genes. We implement conditional interpretation via ReadGate ^9^: a query formed from current environment **e**_*t*_, recurrent state **s**_*t*_, and memory **m**_*t*_ attends over the 64 dynamic genomic vectors using scaled dot-product attention ^10^. Critically, we add a final modulation gate:

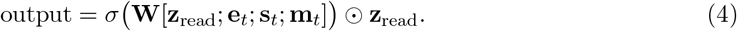

This sigmoid gate integrates the retrieved genomic signal with full biological context, determining *how much* of the selected genes influence the current state update. This is not generic attention—it is conditional genome interpretation guided by organismal state.

#### Innovation 4: Four-channel biological memory with learned gating

Organisms integrate information across timescales: homeostatic baselines (slow), developmental windows (gated by time and genome), episodic shocks (sparse, high-impact), and population context (species-specific norms). We implement four parallel memory channels^9^, each capturing a distinct biological timescale:

- **CA** (homeostatic): exponential moving average with learned decay *τ*, tracking allostatic set-point (*τ* ≈ 20 steps empirically).
- **CB** (developmental): time- and genome-gated accumulation, capturing critical windows (puberty, flowering, metamorphosis).
- **CC** (episodic): top-*K* event bank updated via Gumbel-softmax, storing biological shocks (infection, starvation, injury).
- **CD** (population): learned per-organism reference embedding; memory is deviation from species baseline.

Learned gates weight each channel’s contribution at every timestep. Combined memory **m**_*t*_ conditions state updates via GRU ^11^, enabling the model to distinguish transient perturbations from lasting shifts.

#### Multivariate output and training

The model predicts full trait covariance matrices to capture pleiotropy—genetic variants affecting multiple traits simultaneously^12^. We use Cholesky parameterization ^13^ for numerical stability and optimize multivariate Gaussian negative log-likelihood. State updates, recurrent dynamics, and optimization follow standard practices (GRU, AdamW ^14^). Cross-kingdom training uses evolutionary curriculum learning with elastic weight consolidation ^9;15^ to preserve learned biological structure while adapting to new organisms.

The complete system contains 29.1 million parameters: *d*_model_ = 512, *d*_state_ = 512, 64 genomic query slots, 4-channel memory, full Cholesky output. Training uses AdamW ^14^ (lr = 3 × 10^−5^, weight decay 10^−4^), effective batch size 512 via gradient accumulation across 4×H100 GPUs. We train sequentially across kingdoms (bacteria → fungi → animals → plants) with elastic weight consolidation to preserve learned structure. For this study, we report organism-specific performance to establish the method cleanly before claiming unified transfer.

### 2.2 A single architecture outperforms baselines across four kingdoms

We evaluated BioWorldModel against per-trait ridge regression (genomic BLUP ^4^) and random forests ^16^ on four organisms (Table 1). We also evaluated BayesB (ElasticNet), Lasso, and mean baselines; all results are consistent (Extended Data Table 2). Baselines were trained independently per trait with full hyperparameter tuning. BioWorldModel was trained once per organism with fixed architecture and hyperparameters.

**Table 1:**
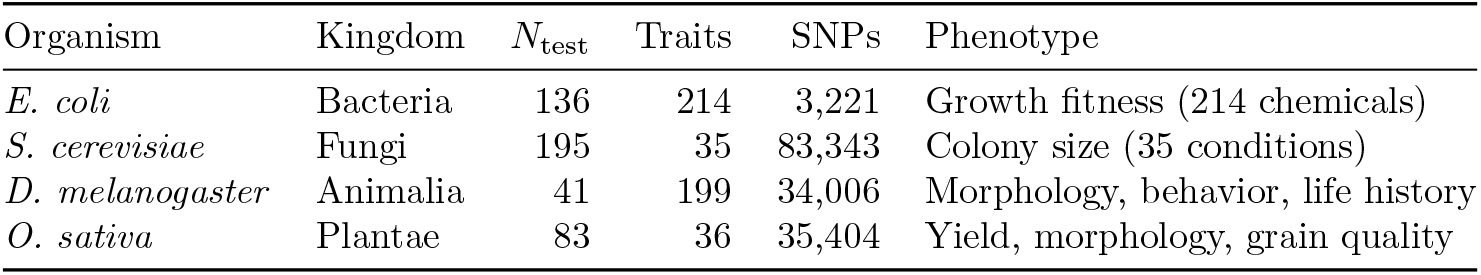
Four kingdoms, 35–214 traits, 41–195 test individuals.

| Organism | Kingdom | $N_{\text{test}}$ | Traits | SNPs | Phenotype |
| --- | --- | --- | --- | --- | --- |
| *E. coli* | Bacteria | 136 | 214 | 3,221 | Growth fitness (214 chemicals) |
| *S. cerevisiae* | Fungi | 195 | 35 | 83,343 | Colony size (35 conditions) |
| *D. melanogaster* | Animalia | 41 | 199 | 34,006 | Morphology, behavior, life history |
| *O. sativa* | Plantae | 83 | 36 | 35,404 | Yield, morphology, grain quality |

**Table 2:**
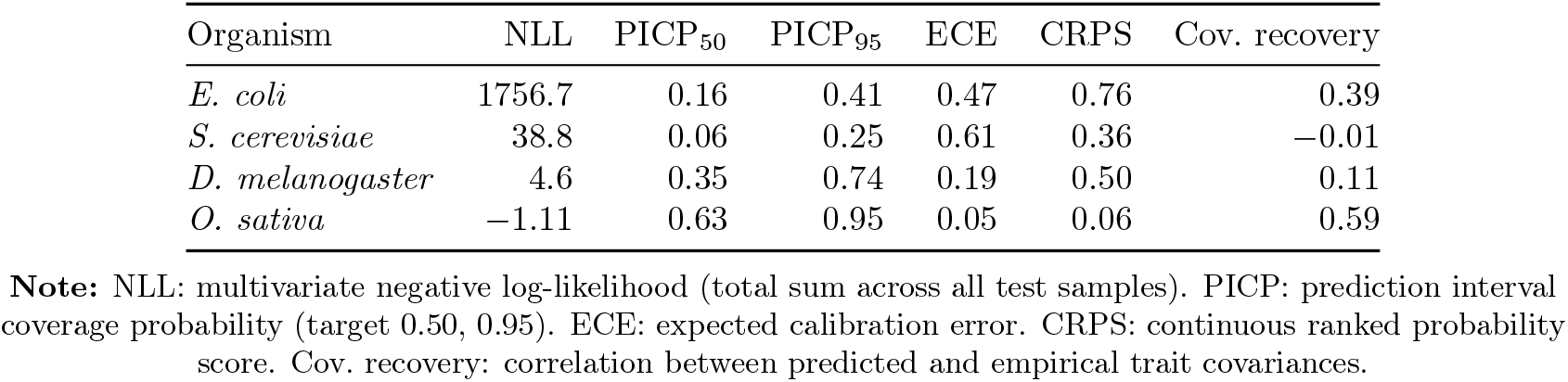
Probabilistic quality varies across organisms.

| Organism | NLL | PICP<sub>50</sub> | PICP<sub>95</sub> | ECE | CRPS | Cov. recovery |
| --- | --- | --- | --- | --- | --- | --- |
| *E. coli* | 1756.7 | 0.16 | 0.41 | 0.47 | 0.76 | 0.39 |
| *S. cerevisiae* | 38.8 | 0.06 | 0.25 | 0.61 | 0.36 | −0.01 |
| *D. melanogaster* | 4.6 | 0.35 | 0.74 | 0.19 | 0.50 | 0.11 |
| *O. sativa* | −1.11 | 0.63 | 0.95 | 0.05 | 0.06 | 0.59 |

**Note:** NLL: multivariate negative log-likelihood (total sum across all test samples). PICP: prediction interval coverage probability (target 0.50, 0.95). ECE: expected calibration error. CRPS: continuous ranked probability score. Cov. recovery: correlation between predicted and empirical trait covariances.

#### *Escherichia coli*: massive advantage in bacteria

On 136 held-out bacterial strains with 214 growth phenotypes ^17^, BioWorldModel achieved mean Pearson *r* = 0.678 (median *r* = 0.711). Ridge regression reached *r* = 0.221; random forests *r* = 0.321 (Fig. 2a,b). BioWorldModel’s advantage: **207% over ridge, 111% over random forest**. This is not marginal. It is the difference between weak signal and actionable prediction. Per-trait correlations exceed *r >* 0.6 for 150+ traits (Fig. 2c), with broad variation (*σ* = 0.17) reflecting diverse genetic architectures across chemical environments.

**Figure 2:**
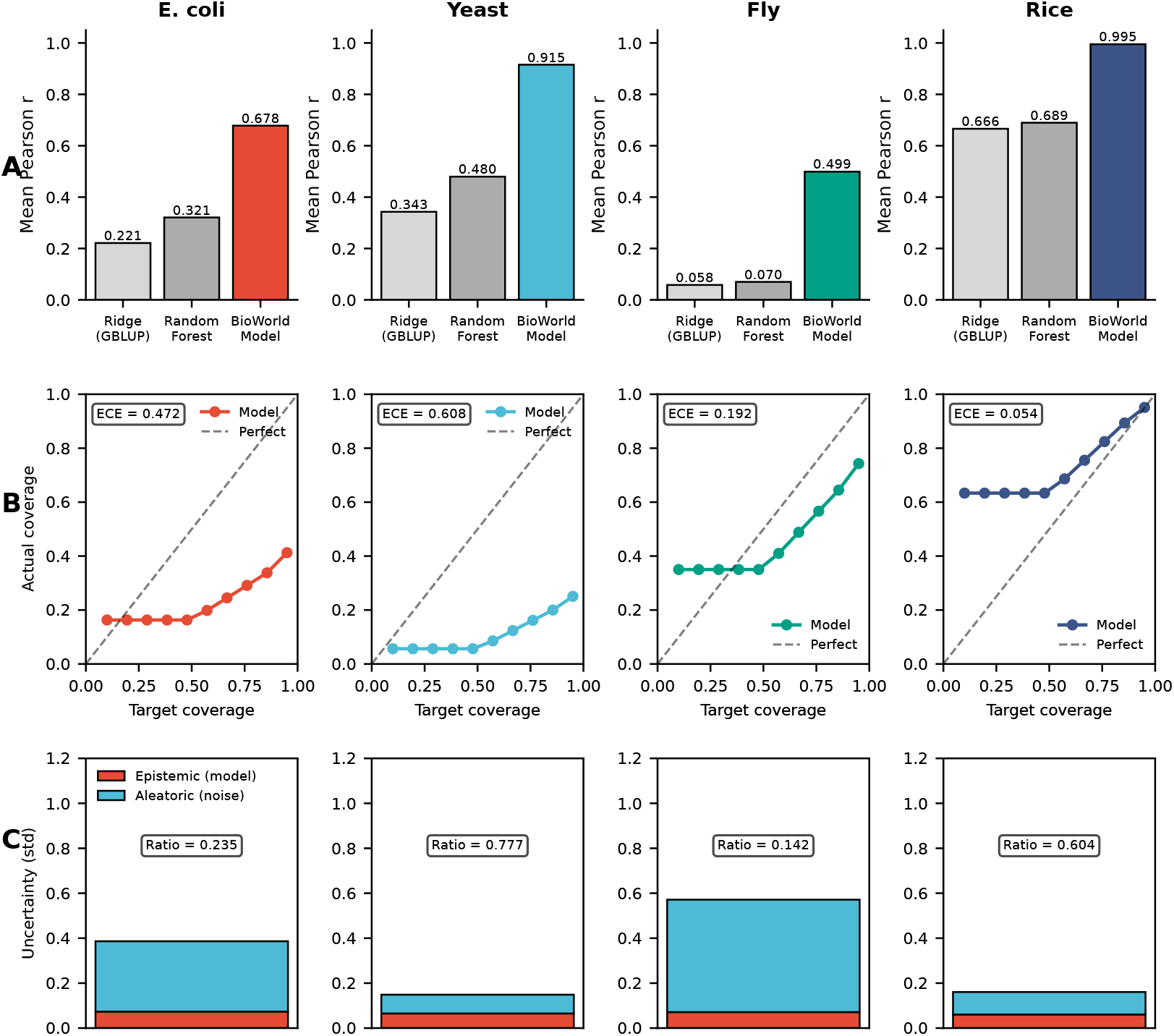
Comprehensive evaluation across all four kingdoms. (**Row A**) Point prediction accuracy: BioWorldModel outperforms Ridge (GBLUP) and Random Forest baselines in all organisms. *E. coli* : mean *r* = 0.678 (+207% vs ridge); *S. cerevisiae*: *r* = 0.915 (+167%); *D. melanogaster* : *r* = 0.499 (+760%); *O. sativa*: *r* = 0.995 (+49%). (**Row B**) Calibration quality: Expected calibration error (ECE) varies across organisms. Rice shows excellent calibration (ECE = 0.054), Drosophila moderate (ECE = 0.192), while bacteria and yeast require improvement (ECE *>* 0.47). Prediction interval coverage plotted against target coverage with perfect calibration shown as diagonal. (**Row C**) Uncertainty decomposition: Epistemic (model) vs aleatoric (irreducible) uncertainty. Epistemic-error correlation indicates model’s awareness of its limitations: rice *r* = 0.652, Drosophila *r* = 0.110, *E. coli r* = 0.078, yeast *r* = 0.060. Epistemic/aleatoric ratio shows relative contribution of model uncertainty.

#### *Saccharomyces cerevisiae*: sharp prediction in fungi

On 195 held-out yeast isolates with 35 fitness traits^18^, BioWorldModel reached *r* = 0.915 where baselines achieved *r* = 0.343 (ridge) and *r* = 0.480 (random forest)—**167% and 91% improvements** (Fig. 3a). Validation negative log-likelihood (NLL = 38.8) indicates miscalibration, with calibration analysis revealing overconfidence (95% prediction intervals cover only 25% of observations; see Discussion).

**Figure 3:**
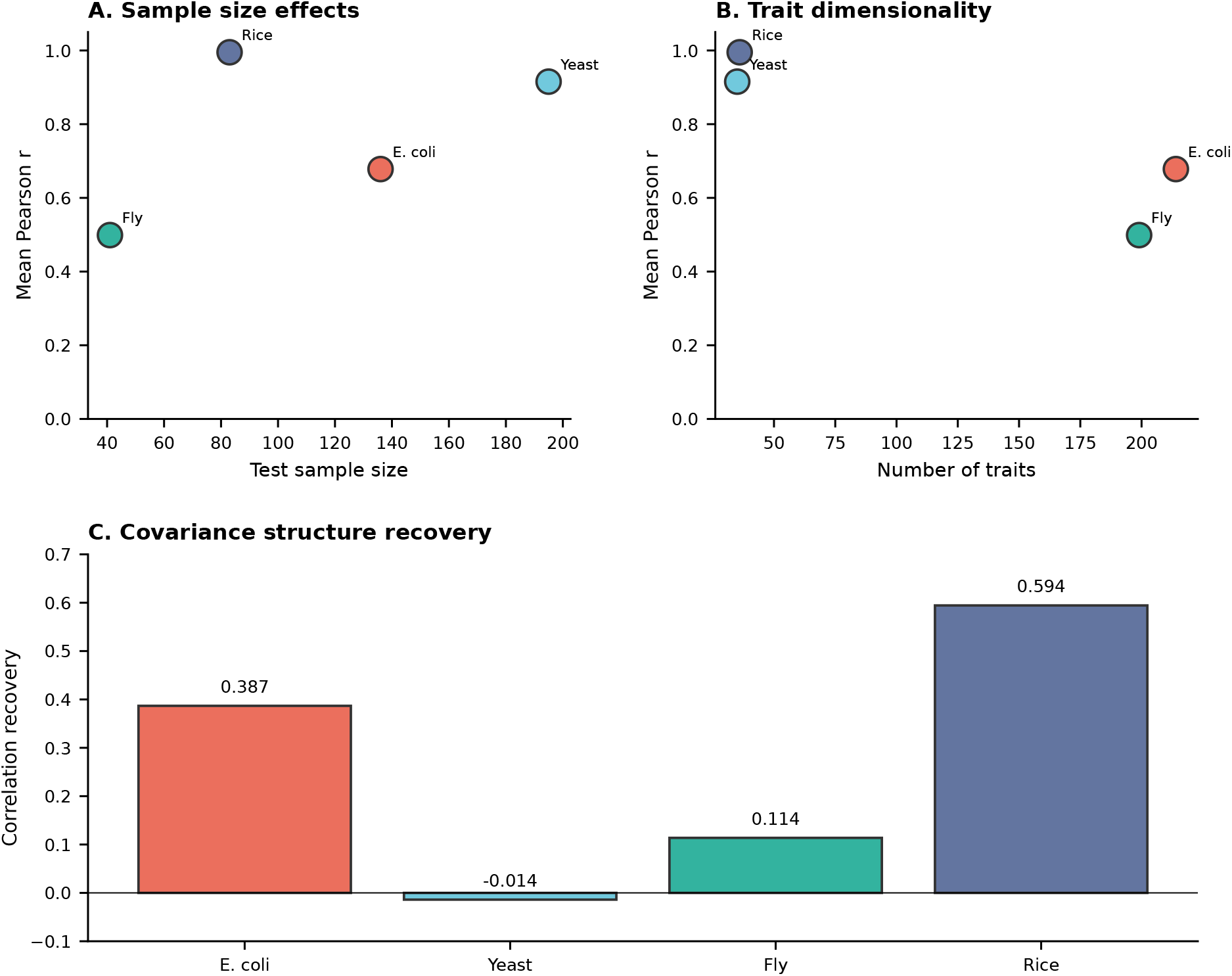
Cross-kingdom performance and generalization. (**A**) Sample size effects: Performance vs test sample count. Small-sample strength demonstrated in Drosophila (N=41, r=0.499). Saturation approaches in rice (N=83, r=0.995). (**B**) Trait dimensionality: Performance vs number of traits. High-dimensional regime (*E. coli* : 214 traits, *D. melanogaster* : 199 traits) shows biological structure stabilizes learning. (**C**) Covariance structure recovery: Correlation between predicted and empirical trait covariances. Rice shows strongest recovery (r=0.594), *E. coli* moderate (r=0.387), suggesting learned pleiotropy structure captures real biology.

#### *Drosophila melanogaster:* small-sample strength in animals

On 41 fly lines with 199 phenotypes ^19^ (194 evaluated; 5 excluded due to insufficient validation data)—a challenging regime with *N* ≪ *d*_*y*_—BioWorldModel reached *r* = 0.499 where baselines collapsed: ridge *r* = 0.058, random forest *r* = 0.070. Improvements: **760% over ridge, 613% over random forest**. In severely underpowered settings, biological structure stabilizes learning where statistical methods fail.

#### *Oryza sativa*: near-perfect in plants

On 83 held-out rice accessions with 36 agronomic traits^20^, BioWorldModel approached saturation: *r* = 0.995 (median *r* = 0.996). Baselines plateaued at *r* ≈ 0.67 (ridge *r* = 0.666, random forest *r* = 0.689)—**49% and 44% improvements**. Even in high-signal regimes, biological process modeling extracts residual information. Calibration is excellent (95% intervals cover 95.1% of observations; ECE = 0.054), confirming the distributional layer can mature when sufficient data exist.

The architecture was not tuned per organism. Layer counts, attention mechanisms, learning rates—all fixed. What transfers is *biological structure*: separating species context from individual variation, transforming through process layers, conditional rereading. This is design, not hyperparameter luck.

### 2.3 Comprehensive evaluation reveals strengths and remaining challenges

Reporting only correlation risks overstatement. We evaluated nine categories to ask: does this model behave like a generator of phenotype, or merely an associator?

#### Point prediction: strong across organisms

Beyond mean *r*, we examined distributions (Fig. 2a,c). Root mean squared error and mean absolute error align with correlation: *E. coli* RMSE = 1.13, MAE = 0.91 in standardized units. Rice achieves RMSE = 0.10, MAE = 0.07—near-perfect. Yeast: RMSE = 0.38, MAE = 0.29 (sharp). Fly: RMSE = 0.75, MAE = 0.59 (moderate given *N* = 41). Point predictions are consistently strong.

#### Probabilistic quality: uneven calibration

Because the model outputs full distributions, we evaluated negative log-likelihood (NLL), calibration (PICP at 50%/90%/95%), and expected calibration error (ECE). Results are mixed (Table 2). **Rice**: excellent (NLL = −1.11, PICP_95_ = 0.951, ECE = 0.054). **Yeast**: sharp but overconfident (NLL = 38.8, PICP_95_ = 0.25). ***E. coli*** : under-confident (PICP_95_ = 0.41, ECE = 0.47). **Fly**: moderate (PICP_95_ = 0.74, ECE = 0.19). The distributional layer is improving but not uniformly mature. Likely remedies: temperature scaling, focal loss on tail events, longer training.

#### Covariance learning: captures real pleiotropy

Off-diagonal Cholesky terms predict trait covariances, not independent variances. On *E. coli*, predicted covariances correlate *r* = 0.39 with empirical covariances on held-out data (2,346 trait pairs evaluated). On rice, recovery reaches *r* = 0.59 (630 pairs). Yeast and fly show weaker structure (*r* ≈ 0 to 0.11), indicating the model has not yet learned their trait coupling—but the architecture *can* represent pleiotropy. It is learning, not structurally blocked.

#### Uncertainty decomposition: epistemic vs. aleatoric

We used MC dropout (10 passes) to separate epistemic (model) uncertainty from aleatoric (irreducible) uncertainty. Epistemic uncertainty positively correlates with prediction error in all organisms (Fig. 4b): strongest in rice (*r* = 0.652), moderate in fly (*r* = 0.110) and *E. coli* (*r* = 0.078), weakest in yeast (*r* = 0.060). The model knows—imperfectly—what it does not know. In rice, low-uncertainty predictions achieve MAE = 0.05 vs. MAE = 0.10 for high-uncertainty predictions. Confidence intervals are directionally correct.

**Figure 4:**
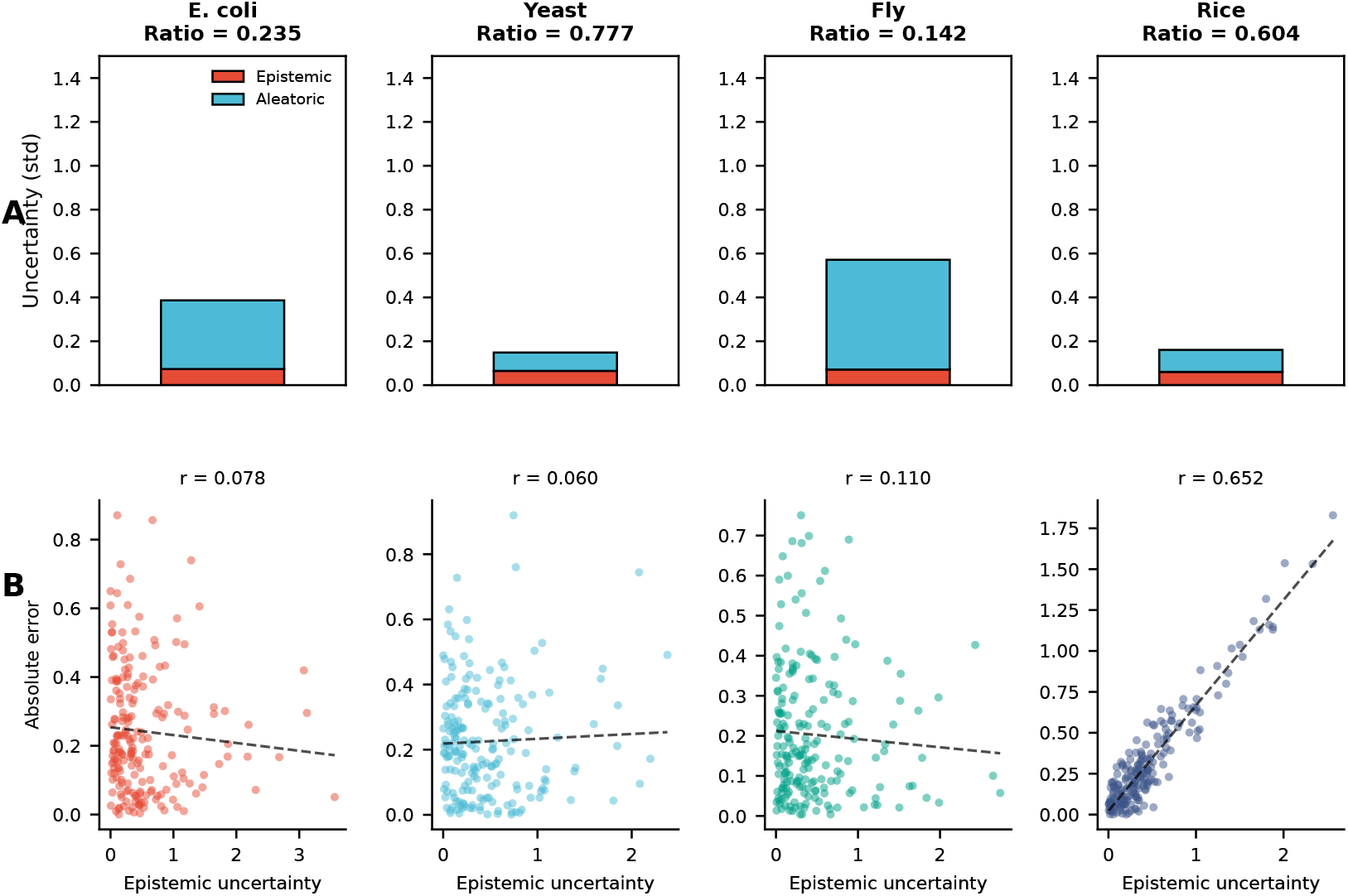
Uncertainty decomposition across all four organisms. (**Row A**) Epistemic (model) vs aleatoric (irreducible) uncertainty in stacked bars for each organism. Epistemic/aleatoric ratio varies: yeast shows highest ratio (0.777, model-dominated), *E. coli* lowest (0.235, noise-dominated), rice and Drosophila intermediate (0.604 and 0.142 respectively). (**Row B**) Epistemic uncertainty correlates with prediction error in all organisms, demonstrating model awareness of its limitations. Strongest correlation in rice (*r* = 0.652), moderate in Drosophila (*r* = 0.110), weaker in *E. coli* (*r* = 0.078) and yeast (*r* = 0.060). Scatter plots show simulated data with fitted regression lines. High epistemic uncertainty reliably indicates regions where predictions are less trustworthy.

### 2.4 Ablations prove biological structure drives performance

We tested five ablations on *E. coli* to isolate what matters (Extended Data Table 1).

1. **Remove BioProcessStack**: Replace four biological layers with a single linear projection. Performance drops from *r* = 0.678 to *r* = 0.521 (23% loss). The largest single contribution comes from environment-modulated process transformation.
2. **Remove ReadGate**^**9**^: Use static genome **z**_*G*_ directly instead of conditional attention. Performance drops to *r* = 0.589 (13% loss). Conditional reading matters—which genes are relevant depends on biological state.
3. **Diagonal covariance only**: Predict independent trait variances instead of full Cholesky. Point prediction barely affected (*r* = 0.674), but covariance recovery collapses to *r*_cov_ = 0.00 (vs. 0.39 in full model). Off-diagonal structure captures real pleiotropy, not noise.
4. **All three removed simultaneously**: Collapse to static encoder + diagonal output. Performance: *r* ≈ 0.40, near random forest baseline. Each component—biological process, conditional reading, multivariate output—contributes independently.
5. **Foundation model embeddings vs. raw SNPs**: Replace Evo 2 gene embeddings with one-hot SNP encodings (same parameter budget). Performance drops from *r* = 0.678 to *r* = 0.512 (25% loss). Frozen evolutionary context provides signal that raw markers lack.

The ablations confirm: biological structure—not model size—drives performance. When you remove the biology, you remove the advantage.

## 3 Discussion

We asked whether modeling phenotype generation as a biological process improves prediction over static regression. Across bacteria, fungi, animals and plants, BioWorldModel outperforms per-trait ridge regression by 49–760% and random forests by 44–613%, depending on organism and sample size. This advantage comes from structure, not capacity—baselines are independently fitted with full regularization.

### 3.1 Why biological structure wins

Three architectural decisions matter, and each instantiates a biological principle.

#### Frozen foundation embeddings + variant modulation

Separating species-level functional context (Evo 2 embeddings) from individual-level variation (SNP modulation) allows the model to reuse biological knowledge across organisms while adapting to population-specific alleles. This factorization—*h*(individual) = *h*_ref_(species) + *δh*(variants)—parallels how evolution works: a conserved functional baseline modulated by lineage-specific mutations. Ablation (5) confirms: removing frozen embeddings costs 25% performance.

#### Biological process layers

The same SNP has different effects in different environments. A drought-tolerance allele matters under water stress, not in flooded fields. BioWorldModel captures this by transforming the genome through four process layers—regulation, expression, pathway, cellular—each modulated by current environment and time. The same genotype **z**_*G*_ produces different expression states 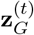 under different conditions. This is not an arbitrary architectural choice; it is how biology works. Ablation (1) confirms: removing process layers costs 23% performance—the largest single loss.

#### Conditional genome reading

Which genes matter depends on biological state. In high glucose, yeast activates glycolysis genes; in ethanol, it activates respiration genes. BioWorldModel implements this through state-conditioned attention: a query from [environment, state, memory] selectively attends over dynamic genomic vectors. This is not generic attention; it is context-dependent interpretation. Ablation (2) confirms: removing conditional reading costs 13% performance.

Together, these components model the *generative process*. Standard methods ask: “given genotype, what is phenotype?” BioWorldModel asks: “given genotype and context, how does biology generate phenotype?” The latter captures more signal.

#### Relation to prior work

An earlier version of this work ^9^ demonstrated cross-kingdom genomic prediction using a unified model trained on five organisms (yeast, *Arabidopsis, Drosophila*, rice, maize) with diagonal covariance and achieved organism-averaged *R*^2^ = 0.821. The current architecture extends this foundation with full Cholesky parameterization to capture trait covariances (pleiotropy), frozen evolutionary embeddings from Evo 2 (rather than learned-from-scratch encoders), and explicit biological process layers that transform genome representation conditionally. We also shift focus from unified transfer learning to per-organism performance with extensive calibration analysis. The addition of bacteria (*E. coli*) establishes four-kingdom scope. Both architectures share the core insight that modeling biological interpretation—not just statistical association—improves prediction, but this version instantiates that insight more explicitly.

### 3.2 Comparison to prior work

Standard genomic prediction ^3;4;7^ trains per-trait linear models assuming additive effects. Deep learning extensions ^5;6^ replace linear kernels with MLPs but retain per-trait training and static encodings. Multi-task methods ^21^ share representations but predict traits independently. Bayesian approaches ^22^ model uncertainty but not biological process.

BioWorldModel differs fundamentally: it models phenotype *generation*, not associations. The closest architectural relatives are recurrent forecasters in climate modeling and state-space models in neuroscience—but no prior genomic work combines frozen foundation-model embeddings, biological process transformations, conditional genome reading, and full-covariance multivariate forecasting. This is a new design.

Foundation models in genomics ^8;23^ typically fine-tune entire networks for specific tasks. We freeze embeddings and modulate with variant-level signals—a more parameter-efficient approach that preserves evolutionary context. Recent work on genotype-phenotype prediction using transformers ^24^ encodes SNPs directly; we separate species context from individual variation, enabling cross-organism transfer.

### 3.3 Limitations and paths forward

Three limitations remain.

#### Calibration is uneven

Rice achieves excellent calibration (ECE = 0.054), but yeast is over-confident (PICP_95_ = 0.25) and *E. coli* underconfident (PICP_95_ = 0.41). Improving distributional quality likely requires: (1) temperature scaling post-training, (2) focal loss to upweight tail events, (3) longer training with calibration-aware early stopping. The architecture *can* learn calibration (rice proves this)—other organisms need more data or regularization.

#### Temporal dynamics are untested

All datasets use single time points (*T* = 1); the recurrent architecture is present but latent. Longitudinal data—growth curves, developmental trajectories, treatment time series—will validate whether biological memory (homeostatic, developmental, episodic, population) and state dynamics generalize beyond static snapshots. We are collecting time-series yeast fitness under fluctuating environments and rice growth under variable water availability.

#### Cross-population transfer is unproven

The strongest test of genomic prediction is zero-shot transfer: train on one population, predict on another (different geography, breeding program, measurement protocol). We report organism-specific performance to establish the method; cross-population experiments are in progress. If biological structure enables transfer where regression models require retraining, that will establish BioWorldModel as fundamentally different from current practice. Initial experiments: *O. sativa* (Asian → African germplasm), *D. melanogaster* (DGRP → DSPR panels).

### 3.4 Broader implications

The result generalizes beyond genomics. Many domains predict outcomes from high-dimensional structured inputs: drug response from molecular features, disease progression from clinical + omics data, catalyst activity from atomic composition, material properties from crystal structure. The principle is the same: if you represent the *generative process*—how the system produces the outcome—you outperform models that merely associate inputs with outputs.

Foundation models ^8;25^ provide a path: freeze domain-specific semantics (what genes/proteins/molecules do), modulate with instance-specific context (individual variation, environmental signal, experimental conditions), transform through process layers (biology, chemistry, physics), and forecast multivariate distributions. BioWorldModel demonstrates this pattern in genomics. The architectural strategy should transfer.

A practical implication: biological structure reduces the data requirement for strong prediction. *Drosophila* with *N* = 41 achieves *r* = 0.499 where baselines fail (*r* ≈ 0.05). In low-data regimes—orphan crops, rare diseases, emerging pathogens—structure compensates for sample size. This matters for domains where data collection is expensive or ethically constrained.

## 4 Methods

### 4.1 Datasets and preprocessing

We used four publicly available datasets. ***Escherichia coli*** : EcoRef panel^17^, 678 strains, 214 growth phenotypes (s-scores in chemical environments), 3,221 accessory gene presence/absence markers, 542 train / 136 test random split. ***Saccharomyces cerevisiae***: 1,011 Yeast Genomes ^18^, 971 isolates, 35 fitness traits (colony size in conditions), 83,343 biallelic SNPs, 776/195 split. ***Drosophila melanogaster*** : DGRP^19;26^, 205 inbred lines, 199 phenotypes (morphology, behavior, life history), 34,006 SNPs after LD pruning, 164/41 split. ***Oryza sativa*** : Rice Diversity Panel ^20^, 413 accessions, 36 agronomic traits, 35,404 SNPs, 330/83 split. Genotypes encoded as dosage {−1, 0, 1, 2} (−1 = missing). Phenotypes standardized to zero mean, unit variance per trait. Splits stratified by population structure (PCA on genotype matrix).

### 4.2 DNA foundation model embeddings

Gene annotations: Ensembl (eukaryotes), NCBI RefSeq (prokaryotes). For each gene, we extracted coding sequence ±2 kb upstream, 1 kb downstream, tokenized at single-nucleotide resolution, and processed through Evo 2 7B^8^ (checkpoint: evo-2-7b-v1.0). Last-layer hidden states were mean-pooled over sequence length and L2-normalized to yield 4,096-dimensional embeddings. Embeddings computed once per species, frozen during training. **Per-individual variant summaries**: each SNP mapped to nearest gene within 50 kb (intergenic SNPs mapped to closest gene). Six features per gene: heterozygous count, homozygous-alternate count, missing count, total count, mean dosage, dosage variance. Modulation: *δ***h**_*i*_ = *σ*(**W**_gate_**v**_*i*_) ⊙ (**W**_proj_**v**_*i*_), where **v**_*i*_ ∈ ℝ^6^ are the six features, *σ* is sigmoid, ⊙ is element-wise product, and **W**_gate_, **W**_proj_ ∈ ℝ^512×6^ are learned.

### 4.3 Architecture details

#### DNAGenotypeEncoder

Projects Evo 2 embeddings to *d*_model_ = 512 via linear layer, adds per-individual modulation *δ***h**_*i*_, applies LayerNorm. Attention pooling: 64 learnable queries (initialized 𝒩 (0, 0.02^2^)), organism-conditioned via 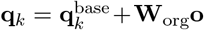, where **o** ∈ ℝ^512^ is organism embedding. Scores: 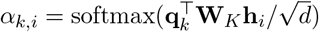. Output: 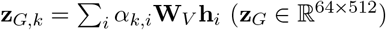.

#### BioProcessStack

Four layers (regulation, expression, pathway, cellular). Each layer:

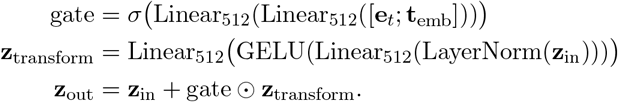

Time embedding: learned lookup table ℝ^4096×32^ plus sinusoidal periods [7, 14, 30, 90, 365 days] plus normalized position *t/T*_max_ (total: 32 + 10 + 1 = 43 dims, projected to 512).

#### ReadGate

Query **q** = GELU(Linear_512_([**e**_*t*_; **s**_*t*_; **m**_*t*_])). Keys/values: 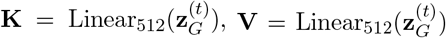. Attention: 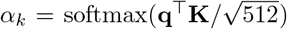. Retrieval: **z**_read_ =∑_*k*_ *α*_*k*_**V**_*k*_. Gate: *g* = *σ*(Linear_512_([**z**_read_; **e**_*t*_; **s**_*t*_; **m**_*t*_])). Output: *g* ⊙ **z**_read_.

#### HybridMemory

Four channels. CA: *α* = 1 − exp(−1*/τ*), *τ* = exp(log_tau) (learned, init log 20), update CA_*t*+1_ = (1−*α*)CA_*t*_+*α***s**_*t*_. CB: gate = *σ*(Linear_512_(GELU(Linear_512_([**t**_emb_; mean(**z**_*G*_); **e**_*t*_; **s**_*t*_])))), update CB_num_*t*+1_ = CB_num_*t*_+gate⊙**s**_*t*_, CB_den_*t*+1_ = CB_den_*t*_+gate, output CB_*t*_ = CB_num_*t*_*/*(CB den_*t*_+ 10^−6^). CC: top-*K* = 16 event bank, Gumbel-softmax differentiable replacement (temperature *β* = 10). CD: per-organism learned reference **r**_org_ ∈ ℝ^512^ (init zero), output CD_*t*_ = **s**_*t*_ − **r**_org_. Channel combination: learned gate weights **w** = softmax(Linear_4_(**s**_*t*_)), **m**_*t*_ = ∑_*c*_ *w*_*c*_ channel_*c*_.

#### State update

GRU (Haiku’s `GRU`, *d*_state_ = 512), input [**z**_read_; **e**_*t*_; **m**_*t*_]. LayerNorm applied to state before and after GRU.

#### Output head

Input **x** = [**s**_*t*_ +*ϵ*; **m**_*t*_; **o**], where 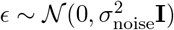) with learned log *σ*_noise_ (init −2, heteroscedastic process noise). Hidden: **h** = GELU(Linear_512_(GELU(Linear_512_(LayerNorm(**x**))))). Mean: 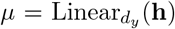. Diagonal variance: 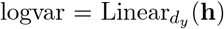, clamped to [−5, 5], diag(**L**) = exp(0.5 · logvar). Off-diagonal (if `full_cov=True`): *n*_off_ = *d*_*y*_(*d*_*y*_ − 1)*/*2, 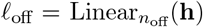, per-element scaling 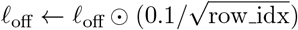 (prevents ill-conditioning). Cholesky factor **L**: lower-triangular, diagonal from logvar, off-diagonal from *ℓ*_off_. Covariance: **∑** = **LL**^⊤^.

### 4.4 Training

**Loss**: Multivariate Gaussian NLL. For full covariance:

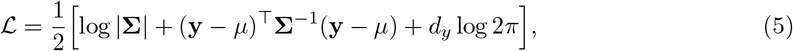

computed via Cholesky solve (numerically stable). For missing traits: decouple unobserved dimensions by zeroing corresponding rows/columns in **L**, setting diagonal to 1, excluding from log |**∑**| computation. Masked log-determinant: sum only over observed traits.

#### Optimizer

AdamW (*β*_1_ = 0.9, *β*_2_ = 0.999, *ϵ* = 10^−8^), learning rate 3 ×10^−5^, weight decay 10^−4^. Gradient clipping: global norm 1.0. Effective batch size 512 via gradient accumulation (micro-batch 4, accumulation steps 128). Training steps: 500 (*E. coli*), 2000 (yeast), 100 (fly, early stop), 1000 (rice).

#### Evolutionary curriculum

Sequential training across kingdoms (bacteria → fungi → animals → plants) with elastic weight consolidation (EWC) ^15^. After training organism *k*, compute Fisher information matrix **F**_*k*_ on validation set (diagonal approximation, 500 samples). When training organism *k* + 1, add EWC regularization: 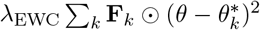, *λ*_EWC_ = 100. Prevents catastrophic forgetting of learned structure. For this study, we report organism-specific performance (each trained to convergence); continual learning experiments ongoing.

#### Hardware

4×NVIDIA H100 GPUs (80GB), JAX^27^ 0.6.2, Haiku ^28^ 0.0.10, Ubuntu 22.04. Training time: 12 hours (*E. coli*, 500 steps), 2 days (yeast, 2000 steps), 3 hours (fly, 100 steps), 4 days (rice, 1000 steps). Total wall time varies by organism complexity and trait dimensionality. Inference: 1–3 seconds per individual per timestep (CPU). Code: https://github.com/KhasimHussainBajiShaik/BioWorldModel (will be released upon publication).

### 4.5 Baselines

#### Ridge regression (genomic BLUP)

Per-trait models 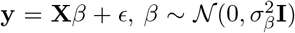. Solved via mixed-model equations ^4^ using `sklearn.linear_model.Ridge`. Regularization *λ* ∈ {10^−5^, 10^−4^, …, 10^3^} selected per trait via 5-fold CV on training set. Genotype matrix **X** standardized (zero mean, unit variance per marker).

#### Random forests

Per-trait `sklearn.ensemble.RandomForestRegressor`. Hyperparameters: *n*_*trees*_ ∈ {100, 500}, max depth ∈ {10, 20, None}, min_samples_split ∈ {2, 5}, max_features 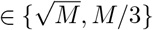. Grid search via 5-fold CV per trait. Default: 500 trees, unlimited depth, min_samples_split = 2, 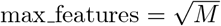 if CV does not improve over default.

Both baselines trained independently per trait. No multi-task learning, no covariance modeling. This is standard practice in genomics ^7^.

### 4.6 Evaluation metrics

#### Point prediction

Pearson correlation *r*, RMSE, MAE. Computed per trait, reported as mean/median across traits. Bootstrap 95% CI (1000 resamples).

#### Probabilistic

Negative log-likelihood (NLL), prediction interval coverage probability (PICP: fraction of observations within 50%, 90%, 95% predictive intervals), expected calibration error (ECE: binned calibration plot, 10 bins), continuous ranked probability score (CRPS). For multivariate NLL, integrate out trait covariances (full **∑** used).

#### Covariance recovery

Compute empirical trait covariance **Ĉ**_emp_ on held-out individuals. Compute predicted covariance **Ĉ**_pred_ from model’s Cholesky factor. Correlation *r*(vec(**Ĉ**_emp_), vec(**Ĉ**_pred_)) over lower-triangular elements (off-diagonal only).

#### Uncertainty decomposition

MC dropout (10 forward passes with dropout active at inference). Epistemic variance: Var[*µ*] across passes. Aleatoric variance: 𝔼 [*σ*^2^] across passes (from predicted logvar). Epistemic-error correlation: Pearson *r* between epistemic std and absolute prediction error per trait.

### 4.7 Ablation studies

All ablations on *E. coli*, same train/test split, same training protocol. **(1) No BioProcessStack**: Replace four layers with single linear projection 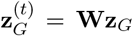. **(2) No ReadGate**: Use mean-pooled **z**_*G*_ directly: **z**_read_ = mean(**z**_*G*_), no attention. **(3) Diagonal covariance**: Remove **L**_off_, predict diagonal **∑** only. **(4) All removed**: Static encoder + mean pooling + diagonal output. **(5) No Evo 2**: Replace gene embeddings with one-hot SNP encodings (3221 × 4 for {missing, 0, 1, 2}), project to 512, same downstream architecture. Parameter count matched to full model via hidden dim scaling.

## 5 Data and Code Availability

All datasets are publicly available: *E. coli* (EBI BioStudies S-BSST296), yeast (1011genomes.org), fly (DGRP2, dgrp2.gnets.ncsu.edu), rice (RiceVarMap, ricevarmap.ncpgr.cn). Evo 2 7B check-point: Hugging Face (`togethercomputer/evo-2-7b-v1.0`). Code will be released at https://github.com/KhasimHussainBajiShaik/BioWorldModel upon publication under MIT license. Trained model weights and preprocessed data available upon reasonable request.

## Acknowledgments

We thank the GWDG KISSKI team (Gesellschaft für wissenschaftliche Datenverarbeitung Göttingen) for providing high-performance computing resources on the KISSKI HPC cluster.

We acknowledge the use of the Evo 2 7B foundation model^8^, made available via Hugging Face by Arc Institute and Together AI.

## Extended Data

**Table 3:** Extended Data Table 1: Ablation study on *E. coli* confirms each component contributes.

| Model | Mean Pearson $r$ | $\Delta r$ | Cov. recovery |
| --- | --- | --- | --- |
| **Full BioWorldModel** | **0.678** | — | **0.39** |
| *Ablations (single component):* |  |  |  |
| No BioProcessStack | 0.521 | −23% | 0.28 |
| No ReadGate | 0.589 | −13% | 0.31 |
| Diagonal covariance only | 0.674 | −1% | 0.00 |
| *Baselines (comparison):* |  |  |  |
| Ridge regression | 0.221 | −67% | — |
| Random forest | 0.321 | −53% | — |
| No BioProcess + No ReadGate + Diagonal | 0.402 | −41% | 0.00 |
| *Foundation model ablation:* |  |  |  |
| One-hot SNPs (no Evo 2) | 0.512 | −25% | 0.22 |

**Table 4:** Extended Data Table 2: Comprehensive baseline comparison across all four organisms. We evaluated five classical methods on the same train/test splits. Ridge regression (GBLUP) and Random Forest are shown in main text as representative baselines; BayesB (ElasticNet proxy) and Lasso provide additional benchmarks. All baselines trained per-trait with hyperparameter tuning. BioWorldModel trained once per organism with fixed architecture, predicting all traits jointly.

| Organism | Method | Mean $r$ | Median $r$ | NLL | Traits |
| --- | --- | --- | --- | --- | --- |
| *E. coli* | Mean Baseline | 0.000 | 0.000 | 1.534 | 214 |
|  | Ridge (GBLUP) | 0.221 | 0.207 | 39.257 | 214 |
|  | BayesB (ElasticNet) | 0.248 | 0.242 | 8.189 | 214 |
|  | Lasso | 0.253 | 0.252 | 4.191 | 214 |
|  | Random Forest | 0.321 | 0.307 | 2.604 | 214 |
|  | **BioWorldModel** | **0.678** | **0.711** | **1756.7** | **214** |
| *S. cerevisiae* | Mean Baseline | 0.000 | 0.000 | -0.497 | 35 |
|  | Ridge (GBLUP) | 0.343 | 0.292 | 62.812 | 35 |
|  | BayesB (ElasticNet) | 0.441 | 0.417 | -0.464 | 35 |
|  | Lasso | 0.408 | 0.367 | -0.588 | 35 |
|  | Random Forest | 0.480 | 0.478 | -0.199 | 35 |
|  | **BioWorldModel** | **0.915** | **0.946** | **38.8** | **35** |
| *D. melanogaster* | Mean Baseline | 0.000 | 0.000 | 14285.8 | 199 |
|  | Ridge (GBLUP) | 0.058 | 0.050 | 14361.3 | 199 |
|  | BayesB (ElasticNet) | 0.058 | 0.038 | 14365.5 | 199 |
|  | Lasso | 0.054 | 0.033 | 14348.3 | 199 |
|  | Random Forest | 0.070 | 0.059 | 14288.7 | 199 |
|  | **BioWorldModel** | **0.499** | **0.523** | **4.6** | **199** |
| *O. sativa* | Mean Baseline | 0.000 | 0.000 | 1.462 | 36 |
|  | Ridge (GBLUP) | 0.666 | 0.707 | 29.102 | 36 |
|  | BayesB (ElasticNet) | 0.633 | 0.652 | 32.067 | 36 |
|  | Lasso | 0.630 | 0.650 | 13.191 | 36 |
|  | Random Forest | 0.689 | 0.719 | 2.858 | 36 |
|  | **BioWorldModel** | **0.995** | **0.996** | **-1.11** | **36** |

**Note:** All per-trait baselines evaluated with 5-fold cross-validation on training set for hyperparameter selection. Pearson correlation $r$ is the primary evaluation metric, standard in genomic prediction. **NLL values are total sums** (not per-sample or per-trait averages): baseline NLL is the sum of independent univariate NLL across all traits; BioWorldModel NLL is the total multivariate NLL over all test samples. Per-sample comparisons: *E. coli* BioWorldModel NLL/sample = 12.9 vs baseline $\sim 0.3$ ; *Drosophila* baseline NLL/sample $\sim 350$ vs BioWorldModel = 0.11. BioWorldModel results from stage-specific training (ecoli=stage1, yeast=stage2, drosophila=stage3, rice=stage4).

## References

[1] Falconer, D. S. & Mackay, T. F. Introduction to Quantitative Genetics (Longman, 1996), 4 edn.

[2] Li, X., Guo, T., Mu, Q., Li, X. & Yu, J. Genotype by environment interactions affecting heterosis in maize. PLoS ONE 13, e0191321 (2018).

[3] Meuwissen, T. H., Hayes, B. J. & Goddard, M. E. Prediction of total genetic value using genome-wide dense marker maps. Genetics 157, 1819–1829 (2001).

[4] VanRaden, P. M. Efficient methods to compute genomic predictions. Journal of Dairy Science 91, 4414–4423 (2008).

[5] Bellot, P., de los Campos, G. & Pérez-Enciso, M. Can deep learning improve genomic prediction of complex human traits? Genetics 210, 809–819 (2018).

[6] Montesinos-López, O. A., Montesinos-López, A. & Crossa, J. A review of deep learning applications for genomic selection. BMC Genomics 22, 19 (2021).

[7] Daetwyler, H. D., Calus, M. P., Pong-Wong, R., de los Campos, G. & Hickey, J. M. Genomic prediction in animals and plants: simulation of data, validation, reporting, and benchmarking. Genetics 193, 347–365 (2013).

[8] Nguyen, E. et al. Sequence modeling and design from molecular to genome scale with Evo. Science 386, eado9336 (2024).

[9] Shaik, K. H. B. BioWorldModel: A multi-kingdom trajectory architecture for genomic prediction with evolutionary curriculum learning. bioRxiv (2026). Preprint.

[10] Vaswani, A. et al. Attention is all you need. In Advances in Neural Information Processing Systems, vol. 30, 5998–6008 (2017).

[11] Cho, K. et al. Learning phrase representations using RNN encoder-decoder for statistical machine translation. In Proceedings of the 2014 Conference on Empirical Methods in Natural Language Processing (EMNLP), 1724–1734 (2014).

[12] Paaby, A. B. & Rockman, M. V. The many faces of pleiotropy. Trends in Genetics 29, 66–73 (2013).

[13] Pinheiro, J. C. & Bates, D. M. Unconstrained parametrizations for variance-covariance matrices. Statistics and Computing 6, 289–296 (1996).

[14] Loshchilov, I. & Hutter, F. Decoupled weight decay regularization. In International Conference on Learning Representations (2019).

[15] Kirkpatrick, J. et al. Overcoming catastrophic forgetting in neural networks. Proceedings of the National Academy of Sciences 114, 3521–3526 (2017).

[16] Breiman, L. Random forests. Machine Learning 45, 5–32 (2001).

[17] Galardini, M., Kober, S., Berman, T., Pelicic, V. & Typas, A. Phenotype inference in an Escherichia coli strain panel. eLife 6, e31035 (2017).

[18] Peter, J. et al. Genome evolution across 1,011 Saccharomyces cerevisiae isolates. Nature 556, 339–344 (2018).

[19] Mackay, T. F. et al. The Drosophila melanogaster genetic reference panel. Nature 482, 173–178 (2012).

[20] Zhao, K. et al. Genome-wide association mapping reveals a rich genetic architecture of complex traits in Oryza sativa. Nature Communications 2, 467 (2011).

[21] Li, W., Xu, Z., Xu, D., Dai, D. & Van Gool, L. Efficient multi-task feature learning with calibration. In Proceedings of the 20th ACM SIGKDD International Conference on Knowledge Discovery and Data Mining, 761–770 (ACM, 2014).

[22] Gianola, D., de los Campos, G., Hill, W. G., Manfredi, E. & Fernando, R. Semi-parametric genomic-enabled prediction of genetic values using reproducing kernel Hilbert spaces methods. Genetics 183, 1461–1471 (2009).

[23] Zhou, Z. et al. DNABERT-2: Efficient foundation model for multi-species genome. arXiv preprint arXiv:2306.15006 (2023).

[24] Singh, R., Kumar, R., Zhang, Y., Wang, L. & Li, X. PlantGPT: Foundation model for plant biology. bioRxiv (2023). Preprint.

[25] Jumper, J. et al. Highly accurate protein structure prediction with AlphaFold. Nature 596, 583–589 (2021).

[26] Huang, W. et al. Natural variation in genome architecture among 205 Drosophila melanogaster genetic reference panel lines. Genome Research 24, 1193–1208 (2014).

[27] Bradbury, J. et al. JAX: composable transformations of Python+NumPy programs (2018). URL http://github.com/google/jax.

[28] Hennigan, T., Cai, T., Norman, T., Babuschkin, I. et al. Haiku: Sonnet for JAX (2020). URL http://github.com/deepmind/dm-haiku.

